# Structural basis for m²,²G27 formation and tRNA remodeling by human TRMT1L

**DOI:** 10.64898/2026.11.24.768959

**Authors:** Tao Li, Arya Vindu, Katherine Barondeau, Sseu-Pei Hwang, Hui-Chi Tang, Ti-Chun Chao, Catherine Denicourt, Kuang-Lei Tsai

## Abstract

tRNAs acquire complex chemical signatures through ordered maturation pathways in which some enzymes contribute both catalytic and non-catalytic activities. Human TRMT1L installs m²,²G27 in tyrosine tRNA and has been reported to support D-loop acp³U20 formation, but the structural basis connecting these functions is unknown. Here, cryo-electron microscopy and mutagenesis show that TRMT1L is a homodimer that extensively reshapes its tRNA substrate. One protomer clamps the acceptor and T arms and supplies the catalytic pocket, while the other engages the anticodon loop. Charge reversal at these anticodon contacts abolishes methylation without disrupting the dimer, showing that engagement by one protomer is required for catalysis by the other. Cofactor binding is accompanied by reorganization of the D- and anticodon arms that draws G27 into the catalytic pocket. In a second cofactor-bound state an active-site loop (L1) that is disordered without cofactor becomes ordered near the position of the extruded base, suggesting a steric gating mechanism that controls access to the catalytic site. Thus, TRMT1L remodels its substrate rather than closing around it, a property that may explain its requirement for modifications that it does not itself catalyze.

## Introduction

Chemical modifications allow a limited set of RNA building blocks to support remarkably diverse structures and functions ^1^. This chemical diversification is especially extensive in transfer RNAs (tRNAs), where modifications alter base pairing, tertiary folding, stability, and interactions with the translation machinery ^2–4^. Modification levels can change with nutrient availability, oxidative stress, and other physiological cues, altering the functional properties of the tRNA pool and favoring translation of codon-biased mRNAs that support adaptation to these conditions ^5–7^. Dysregulation or loss of tRNA-modifying enzymes is associated with diverse human diseases, underscoring the biological consequences of disrupting this regulation ^8–10^. Defining how these enzymes engage their substrates is therefore essential to understanding how tRNA modification sustains and regulates protein synthesis.

N²,N²-dimethylguanosine (m²,²G) contributes to tRNA tertiary structure by restricting the pairing interactions available to guanosine; at position 26, near the junction of the D arm and anticodon stem, it disfavors competing interactions that can stabilize alternative tRNA conformations ^11–14^. The conserved m²,²G methyltransferase Trm1, represented by TRMT1 in humans, modifies G26 in numerous tRNAs ^15–18^. Its vertebrate paralog, TRMT1-like (TRMT1L), installs m²,²G at the adjacent position 27 exclusively on tRNA-Tyr-GUA ^16,19^. Loss of TRMT1L reduces tRNA-Tyr-GUA abundance, impairs global translation, and increases sensitivity to oxidative stress, establishing its importance for tRNA function and cellular homeostasis ^19^. How two closely related enzymes select different positions and different substrates has remained unclear.

The contribution of TRMT1L to tRNA maturation extends beyond m²,²G27 formation. TRMT1L is also required to maintain normal levels of 3-(3-amino-3-carboxypropyl)uridine at position 20 (acp³U20) on tRNA-Tyr, tRNA-Cys, and tRNA-Ala, a modification installed by the distinct enzyme DTWD1 ^16,19,20^. This requirement is independent of G27 methylation: catalytically inactive TRMT1L restores acp³U20 on tRNA-Cys ^16,19^, while a neurodegeneration-associated TRMT1L variant impairs acp³U modification on selected tRNAs despite retaining G27 methylation ^16^. These separable activities raise a mechanistic question: how does TRMT1L support modification of tRNA by another enzyme without requiring its own methyl-transfer reaction?

A possible explanation is that TRMT1L changes the conformation of its RNA substrates in a manner that facilitates subsequent modification. Fission yeast Trm1 has RNA strand-annealing and dissociation activities and can support tRNA function independently of catalysis, a precedent for chaperone-like activity within the family ^21^. Whether human TRMT1L uses such an activity to support acp³U20 formation is unknown. Addressing this requires knowing how TRMT1L contacts tRNA, whether binding reorganizes the RNA beyond G27, and how any such rearrangement is coupled to G27 engagement.

Here we combine cryo-electron microscopy of human TRMT1L bound to tRNA-Tyr, in cofactor-free (apo) and S-adenosylhomocysteine (SAH)-bound states, with biochemical analysis and structure-guided mutagenesis. TRMT1L is a homodimer in which a single tRNA spans both protomers: one binds the acceptor and T arms and supplies the catalytic pocket, whereas the other engages the anticodon loop. SAH binding is accompanied by reorganization of the D- and anticodon arms and extrusion of G27. In a second SAH-bound state an active-site loop (L1) closes over the pocket while anticodon engagement is weakened, suggesting how the tRNA may be released. Mutagenesis shows that the anticodon contacts, dimer assembly and L1 loop integrity are required for m²,²G27 formation. Together, these structures show how a dimer divides substrate engagement from catalysis, and how cofactor binding accompanies extrusion of the target base.

## RESULTS

### Structure determination of the human TRMT1L-tRNA-Tyr complex

To understand how TRMT1L recognizes tRNA-Tyr and accesses G27, we reconstituted an active TRMT1L-tRNA complex for structure determination. TRMT1L and TRMT1 share a class I *S*-adenosyl-L-methionine (SAM)-dependent methyltransferase fold ^14,22–24^. However, TRMT1L has an N-terminal region of approximately 225 residues, including a predicted zinc-finger domain, that has no counterpart in TRMT1 or in archaeal and bacterial Trm1 enzymes (Fig. 1a). A putative Cys4 metal-binding site, conserved in TRMT1, lies C-terminal to the methyltransferase domain, at the position of the zinc finger in the structure of *A. aeolicus* Trm1 ^24^(Fig. 1a). Full-length human TRMT1L was expressed in insect cells and purified by anti-FLAG affinity chromatography followed by size-exclusion chromatography (SEC) (Supplementary Fig. 1a, b). The SEC profile suggested that recombinant TRMT1L is a homodimer *in vitro*. Co-immunoprecipitation of FLAG-tagged and Myc-tagged TRMT1L from co-transfected HEK293T cells indicated that it also self-associates in cells (Supplementary Fig. 1c). The recombinant protein had methyltransferase activity comparable to TRMT1L purified from HEK 293T cells and formed a stable complex with tRNA-Tyr, as shown by SEC (Fig. 1b and Supplementary Fig. 1d, e). We concentrated the cofactor-free complex to approximately 1 mg/mL, vitrified it and analyzed it by single-particle cryo-electron microscopy (cryo-EM) (Supplementary Fig. 2a).

**Figure 1.**
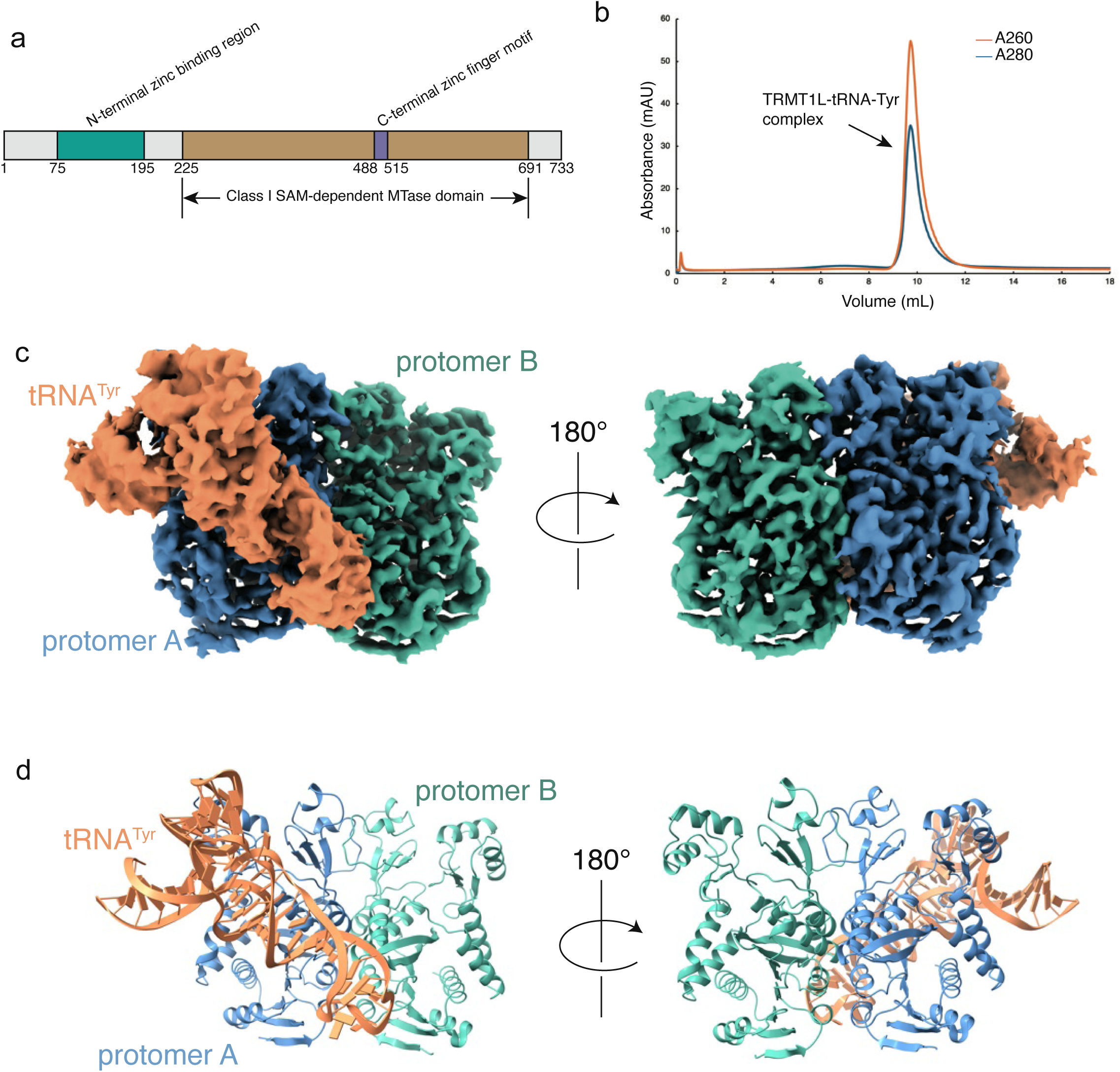
Dimerization and cryo-EM structure of human TRMT1L bound to tRNA-Tyr. **a.** Schematic representation of the domain organization of human TRMT1L, showing the N-terminal zinc-binding region, C-terminal zinc finger motif, and class I SAM-dependent methyltransferase domain. **b.** Size-exclusion chromatography profile of the purified TRMT1L-tRNA-Tyr complex, monitored by absorbance at 260 nm (orange) and 280 nm (blue). The peak corresponding to the TRMT1L-tRNA-Tyr complex is indicated by an arrow. **c.** Cryo-EM density map of the apo TRMT1L-tRNA-Tyr complex shown in two views related by a 180° rotation. TRMT1L protomers A and B are colored blue and teal, respectively, and tRNA-Tyr is colored orange. **d.** Overall atomic model of the apo TRMT1L-tRNA-Tyr complex shown in the same orientations and color scheme as in c.

Two-dimensional class averages showed twofold symmetry of the protein, consistent with a homodimer (Supplementary Fig. 2a). Ab initio 3D reconstruction revealed two populations of homodimers, bound to one or to two tRNA-Tyr molecules. We processed the singly bound complex, the predominant population. After multiple rounds of 3D classification, approximately 60,294 particles yielded a map at an overall resolution of approximately 3.2 Å (Supplementary Fig. 2a). The map showed a single tRNA-Tyr bound to a TRMT1L homodimer (Fig. 1c). We built an atomic model using AlphaFold-predicted structures of TRMT1L and tRNA-Tyr as references (Fig. 1d). Flexible regions, including the N-terminal region and the C-terminal disordered region of TRMT1L, were not modeled because of poorly resolved density.

The homodimer comprises protomers A and B. A single tRNA-Tyr binds primarily to protomer A and makes additional contacts with protomer B (Fig. 1d). The resolved C-terminal region adopts a class I methyltransferase fold with a Rossmann-like α/β architecture and a zinc-finger motif. It provides the platform for dimerization, cofactor binding, tRNA recognition and catalysis. No cofactor density was observed in the catalytic site. L1 (residues 238-250), the loop that precedes the active-site residue Tyr251, is unresolved in this cofactor-free state, and the model of protomer A begins at residue 256. L1 is five residues longer in TRMT1L than in TRMT1, and the corresponding loop in *P. horikoshii* Trm1 is flexible, where it was resolved only in an SAH-bound structure (Supplementary Fig. 3) ^23^.

### Dimerization is important for TRMT1L methyltransferase function

Although *P. horikoshii* Trm1 crystallizes as a dimer, analytical gel filtration indicated that it is monomeric in solution, as reported earlier for *P. furiosus* Trm1 ^23,25^. In contrast, our biochemical and structural analyses demonstrate that human TRMT1L forms a stable and catalytically active homodimer. The two protomers associate through an extensive interface stabilized by three regions of intermolecular interaction (Fig. 2a). At one end, residues D357, H363 and R387 from both protomers contribute to a four-helix bundle (Fig. 2b). In the central region, R414 and R415 of one protomer interact with D476 and N447 of the opposing protomer, forming two symmetric interaction networks (Fig. 2c). At the opposite end, the C-terminal zinc-finger-containing region (residues 484-527) makes additional contacts involving V503, E505, Y508 and H516 (Fig. 2d). Together, these interactions stabilize the homodimer.

**Figure 2.**
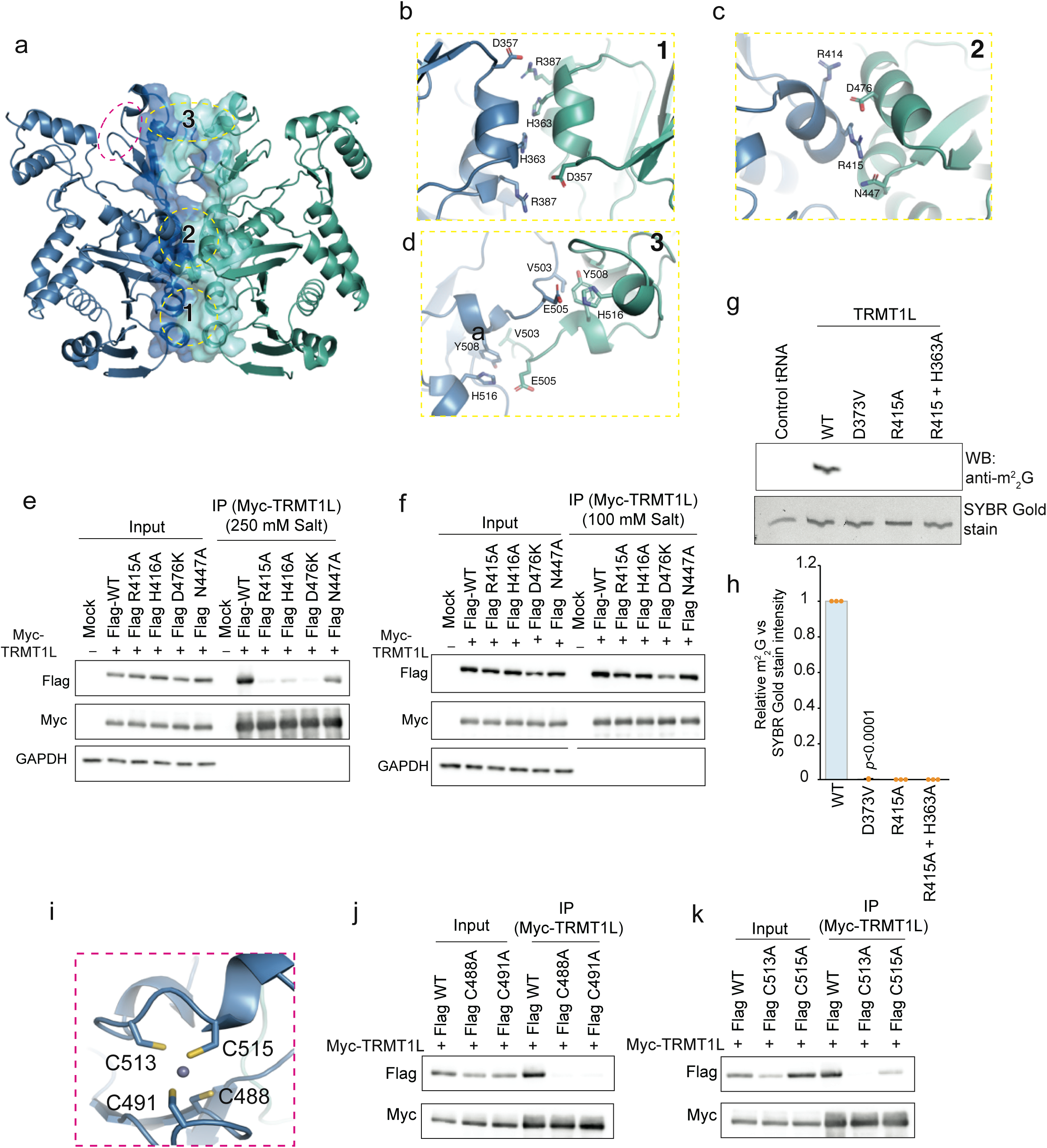
Dimerization is essential for TRMT1L methyltransferase activity. **a.** Overall structure of the TRMT1L homodimer showing the extensive interprotomer interface, highlighted by a semitransparent surface. Three distinct interface regions are indicated by yellow dashed circles (1-3), and the C-terminal zinc finger motif is highlighted by a magenta dashed circle. **b-d.** Close-up views of the three interface regions showing interactions between residues from the two TRMT1L protomers. b, Region 1, involving R387, H363, and D357. c, Region 2, involving R414, R415, N447, and D476. d, Region 3, involving Y508, V503, E505, and H516. **e-f.** Mutations of key residues at the dimer interface disrupt TRMT1L homodimerization. co-IP of Myc-TRMT1L WT with FLAG-tagged TRMT1L WT or the indicated interface mutants expressed in HEK 293T cells and lysed in buffer containing either 250 mM or 100 mM NaCl. **g-h.** Disruption of TRMT1L homodimerization impairs its m²,²G methyltransferase activity. *In vitro* transcribed tRNA-Tyr was incubated with purified Flag-TRMT1L WT, D373V, R415A and R415A/H363A proteins and analyzed by immuno-Northern blotting with m²,²G antibody and densitometry quantification of the m²,²G signals. Data are presented as mean ± SD. N=3. **i.** Close-up view of the C-terminal zinc finger motif, coordinated by C488, C491, C513, and C515, which contributes to TRMT1L homodimerization. **j-k.** Mutations of zinc-coordinating residues impair TRMT1L dimer formation. co-IP of Myc-TRMT1L WT with FLAG-tagged TRMT1L WT or the indicated zinc-coordinating residues expressed in HEK 293T cells and lysed in buffer containing 250 mM NaCl.

To test the functional importance of dimerization, we introduced substitutions at the dimer interface, targeting residues distant from the catalytic site. We co-expressed FLAG-tagged TRMT1L WT or individual interface mutants with Myc-tagged TRMT1L WT and assessed self-association by co-immunoprecipitation (Fig. 2e, f). To distinguish a weakened interaction from a complete loss, we performed the co-immunoprecipitations in lysis buffer containing either 100 mM NaCl, which preserves weaker interactions, or 250 mM NaCl, a more stringent condition. Wild-type TRMT1L self-associated at both salt concentrations, indicating that the dimer is salt-stable. In contrast, R415A, H416A and D476K substantially reduced recovery of Myc-TRMT1L WT, and N447A reduced it, with greater recovery at 100 mM than at 250 mM NaCl (Fig. 2e, f). The substitutions therefore weaken the dimer interface and render it salt-sensitive.

We next asked whether weakening the interface impairs catalysis, using a reconstituted in vitro assay with SAM and in vitro-transcribed tRNA-Tyr ^19,26^. This assay is performed at 50 mM NaCl, because higher salt concentrations reduce TRMT1L activity (Supplementary Fig. 4a). The single interface substitutions that impaired dimerization at 250 mM NaCl retained partial self-association at 100 mM NaCl (Fig. 2e, f), so we also sought a combination that would impair dimer formation further. Double mutants combining R415A with H416, N447 or D476 substitutions yielded no recoverable protein, so we paired R415A with H363A in interface region 1 (Fig. 2b). The R415A/H363A double mutant showed impaired dimer formation at low salt (Supplementary Fig. 4b). Unexpectedly, R415A alone and R415A/H363A both lost m²,²G27 formation relative to wild type (Fig. 2g, h), even though R415A retained partial self-association in cells. Because R415 also contacts the G27-containing region of the tRNA (Fig. 3f), the loss of activity in R415A may reflect weakened dimerization together with altered tRNA engagement.

**Figure 3.**
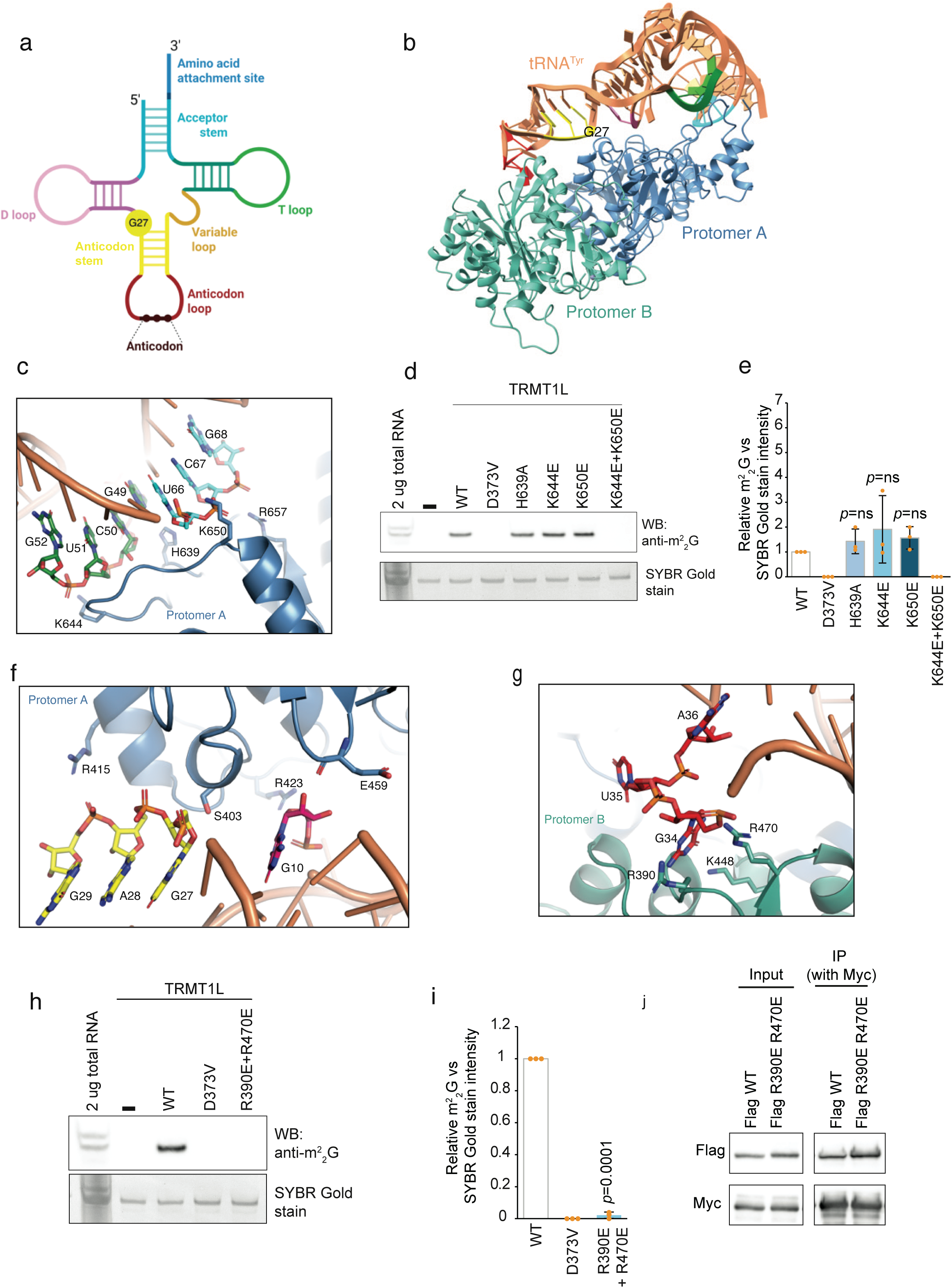
Recognition of tRNA-Tyr by the TRMT1L homodimer. **a.** Schematic representation of the secondary structure of tRNA-Tyr, highlighting its major structural elements and the position of G27. **b.** Overall structure of apo TRMT1L-tRNA-Tyr complex showing how the TRMT1L homodimer recognizes tRNA-Tyr through three distinct interaction regions. TRMT1L protomers A and B are colored blue and teal, respectively, and tRNA-Tyr is colored orange, with the interacting regions highlighted in different colors. **c.** Close-up view of interactions between the C-terminal region of TRMT1L protomer A and the T-stem and acceptor stem of tRNA-Tyr (region I). Key interacting residues are shown as sticks. **d-e.** Mutations of key residues contacting the T-stem impair TRMT1L m²,²G methyltransferase activity. *In vitro* transcribed tRNA-Tyr was incubated with purified Flag-TRMT1L WT, D373V, H639E, K644E, K650E, K644E/K650E proteins and analyzed by immuno-Northern blotting with m²,²G antibody and densitometry quantification of the m²,²G signals. Data are presented as mean ± SD. N=3. **f.** Close-up view of the G27 containing region of tRNA-Tyr recognized by protomer A (region II). G27 remains within the tRNA structure rather than flipping out toward the catalytic pocket in the apo state. **g.** Close-up view of interactions between the anticodon loop of tRNA-Tyr and protomer B (region III). G34 in the anticodon loop interacts with residues including R390, K448 and R470 in protomer B. **h-i.** Mutations of key anticodon loop-interacting residues impair TRMT1L-mediated m²,²G27 formation. *In vitro* transcribed tRNA-Tyr was incubated with purified Flag-TRMT1L WT, D373V, R390E/R470E proteins and analyzed by immuno-Northern blotting with m²,²G antibody and densitometry quantification of the m²,²G signals. Data are presented as mean ± SD. N=3. **j**. Mutations of key anticodon loop-interacting residues does not impair TRMT1L dimer formation. co-IP of Myc-TRMT1L WT with FLAG-tagged TRMT1L WT or the indicated anticodon loop-interacting residues expressed in HEK 293T cells and lysed in buffer containing 250 mM NaCl.

The map also revealed a putative Cys4 zinc-binding site in the C-terminal region of each protomer, formed by Cys488, Cys491, Cys513 and Cys515 (Fig. 2a, i). The geometry is consistent with tetrahedral metal coordination, and a Zn ion was modeled at each site. Structural superposition with *A. aeolicus* Trm1 places the site in the corresponding zinc-binding region ^27^. Although the two zinc-binding sites do not contact each other, they may contribute to the local structural stability required for dimer formation. Consistent with this idea, mutation of the zinc-coordinating residues slightly destabilized TRMT1L and impaired dimer formation, supporting a structural role for this site (Fig. 2j, k). Together, these data indicate that the C-terminal zinc sites help maintain the TRMT1L dimer, which provides the composite surface for tRNA binding.

### Recognition of tRNA-Tyr by TRMT1L

In our structure, tRNA-Tyr engages the TRMT1L homodimer through three distinct regions (Fig. 3a, b). First, the T-stem and acceptor stem interact with the C-terminal region (residues 633-694) of protomer A, where K644, H639, N647, K650 and R667 contact the tRNA backbone (Fig. 3c). This predominantly backbone-directed interaction suggests recognition of the tRNA tertiary fold rather than its sequence. Individual charge-reversal substitutions H639E, K644E and K650E did not reduce m²,²G27 formation relative to wild type (Fig. 3d, e), consistent with binding supported by multiple backbone contacts rather than dependence on any one residue. In contrast, the K644E/K650E double mutant abolishes in vitro m²,²G formation, indicating that these contacts contribute collectively to tRNA engagement (Fig. 3d, e).

Second, the central region of the methyltransferase domain in protomer A contacts the sugar-phosphate backbone of the G27-containing region via S403, R415, R423 and E459, while G27 remains stacked within the tRNA structure (Fig. 3f and Supplementary Fig. 5a, b). Third, the anticodon loop interacts with protomer B, where R390, R470 and K448 contact G34 and the phosphate backbone. No equivalent contacts are observed with protomer A in this state (Fig. 3g).

To test the functional importance of the interactions between protomer B and the anticodon loop, we introduced the double mutant R390E/R470E. Because TRMT1L is a homodimer, these substitutions are present in both protomers; in protomer B, they replace two residues that directly contact G34 and the phosphate backbone with residues expected to disrupt those contacts. The double mutant abolished m²,²G27 formation (Fig. 3h, i) while still assembling as a dimer (Fig. 3j), indicating that the loss of activity reflects the disrupted anticodon contacts rather than a failure of assembly. These results support the structural evidence that both protomers cooperatively engage tRNA-Tyr through a composite substrate-binding interface.

### Cofactor binding remodels the anticodon arm and repositions G27 toward the catalytic pocket

To determine how cofactor binding relates to G27 modification, we added SAH to the TRMT1L-tRNA-Tyr complex before vitrification and determined the structure by cryo-EM (Supplementary Fig. 2b). Three-dimensional classification revealed two conformations of the SAH-bound complex (Supplementary Fig. 2b). In the first, SAH is bound within an elongated, solvent-accessible pocket of the methyltransferase domain (Fig. 4a) and G27 flips out of the tRNA body and projects toward the catalytic pocket while the L1 loop remains disordered. We refer to this as the G27-flipped-out state (Fig. 4b and Supplementary Fig. 5c, d). In the second, SAH binds to the same catalytic pocket. G27 remains stacked in the tRNA body while the L1 loop is stabilized, accompanied by rearrangements in the catalytic core. We refer to this as the G27-unflipped state, described below (Fig. 5a and Supplementary Fig. 5e).

**Figure 4.**
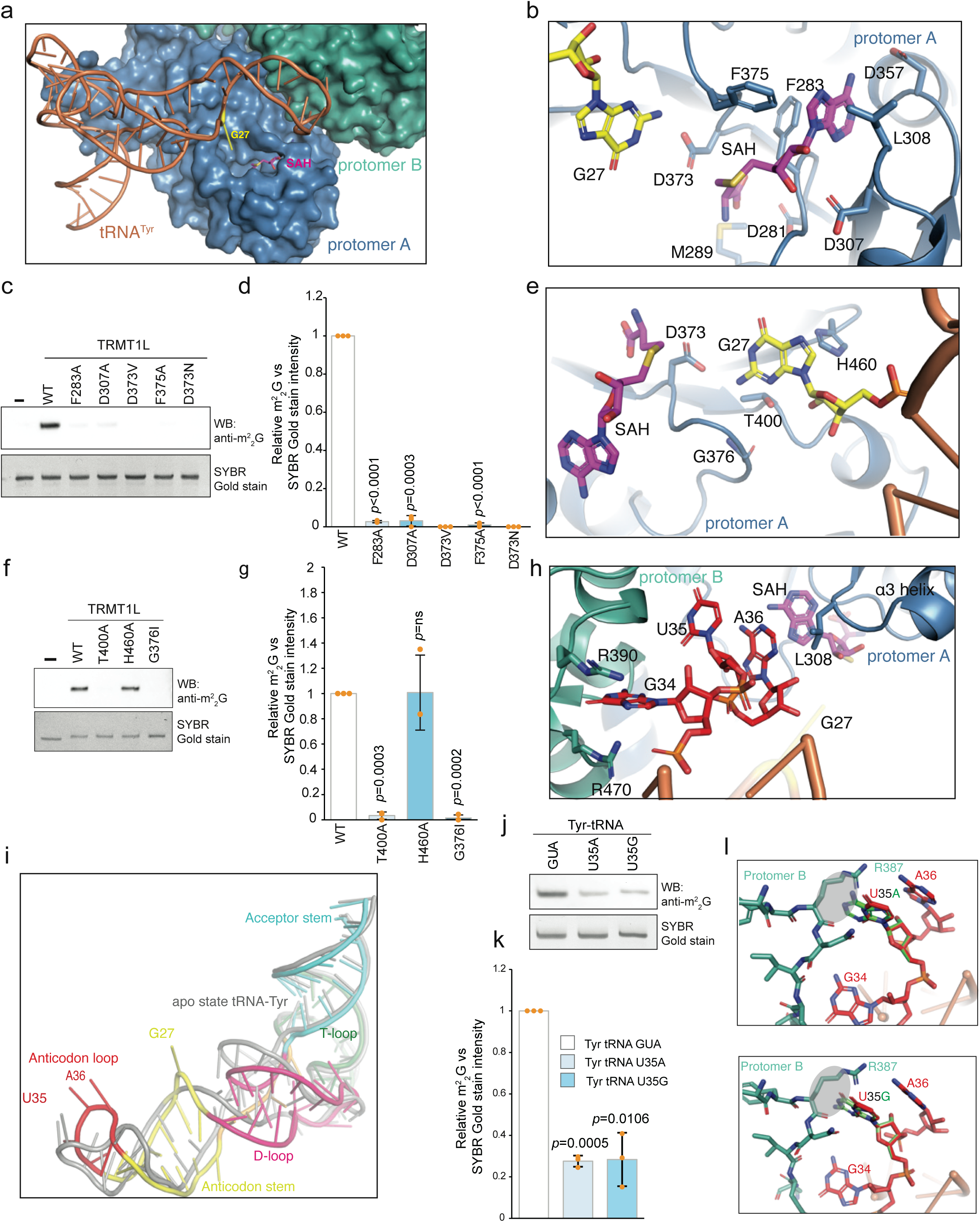
SAH binding induces G27 flipping and conformational changes in tRNA-Tyr. **a.** Overall structure of the SAH-bound G27-flipped-out state of TRMT1L-tRNA-Tyr complex showing SAH bound within a deep catalytic pocket of protomer A. TRMT1L protomers A and B are shown as blue and teal molecular surfaces, respectively, with tRNA-Tyr colored orange. SAH and G27 are highlighted in magenta and yellow, respectively. **b.** Close-up view of the catalytic pocket showing interactions between SAH and surrounding residues, including D281, F283, M289, D307, L308, D373 and F375. **c-d.** Mutations of key SAH-interacting residues abolish TRMT1L-mediated m²,²G27 formation. *In vitro* transcribed tRNA-Tyr was incubated with purified Flag-TRMT1L WT or indicated mutant proteins and analyzed by immuno-Northern blotting with m²,²G antibody and densitometry quantification of the m²,²G signals. Data are presented as mean ± SD. N=3. **e.** Close-up view of the flipped-out G27 in the shallow recess of protomer A, which consists of residues, including D373, G376, T401 and H460. **f-g.** Mutations of key G27-interacting residues abolish TRMT1L-mediated m²,²G27 formation. *In vitro* transcribed tRNA-Tyr was incubated with purified Flag-TRMT1L WT or indicated mutant proteins and analyzed by immuno-Northern blotting with m²,²G antibody and densitometry quantification of the m²,²G signals. Data are presented as mean ± SD. N=3. **h.** Rearrangement of the anticodon loop is stabilized by interactions with both TRMT1L protomers and SAH. G34 interacts with R390 and R470 of protomer B, whereas A36 contacts L308 of protomer A and SAH. **i.** Superimposition of tRNA-Tyr from the apo and SAH-bound G27-flipped-out states reveals conformational rearrangements of the anticodon loop, D-loop and G27 region, while the overall architectures of the acceptor stem and T stem are largely maintained. The apo sate tRNA-Tyr is shown in gray, and the other state tRNA-Tyr is colored by its structural elements. **j-k.** Mutations of U35 in the anticodon of tRNA-Tyr abolish TRMT1L-mediated m²,²G27 formation. *In vitro* transcribed tRNA-Tyr GUA (U35), GAA (U35A) or GGA (U35G) was incubated with purified Flag-TRMT1L WT and analyzed by immuno-Northern blotting with m²,²G antibody and densitometry quantification of the m²,²G signals. Data are presented as mean ± SD. N=3. **l.** Structural models illustrating potential steric clashes upon replacement of U35 with adenine (top) and guanine (bottom) in the anticodon loop of tRNA-Tyr. Both substitutions are predicted to introduce steric clashes with R387 of TRMT1L protomer B, as indicated by the gray ovals. TRMT1L protomer B is shown in teal, with the anticodon loop colored red and the substituted adenine and guanine bases highlighted in green.

**Figure 5.**
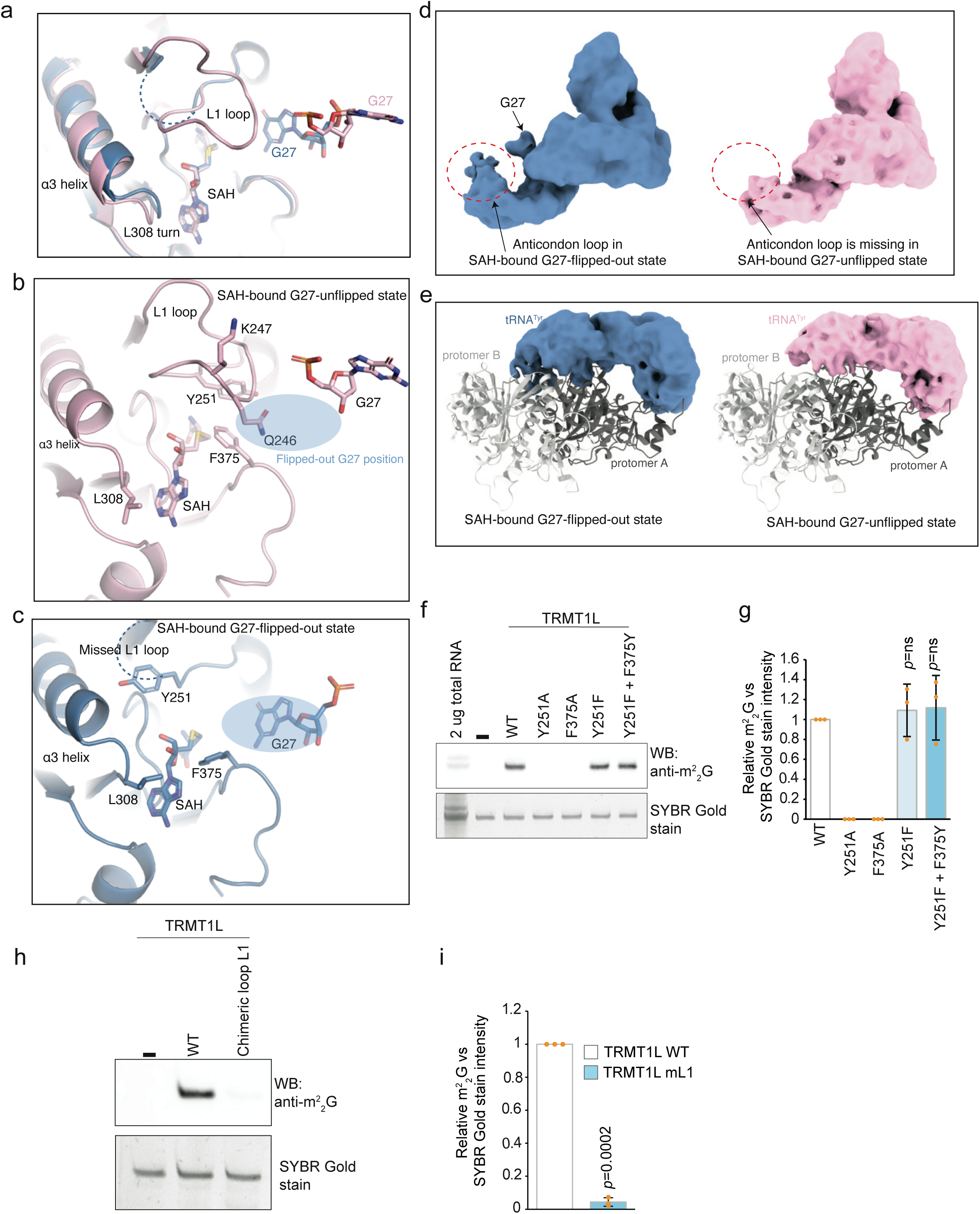
L1 loop stabilization promotes catalytic pocket closure and G27 repositioning. **a.** Superimposition of the two SAH-bound TRMT1L-tRNA-Tyr structures showing conformational rearrangements of the L1 loop, the L308 containing turn and α3 helix, accompanied by repositioning of G27. The G27-flipped-out and G27-unflipped states are shown in blue and pink, respectively. **b.** Close-up view of the catalytic pocket in the SAH-bound G27 unflipped state. Stabilization of the L1 loop is accompanied by rearrangements of Y251 and F375, which contribute to catalytic pocket closure, and displacement of the L308-containing turn and α3 helix. In this state, G27 returns toward its original position within the tRNA structure, while Q246 occupies a position near the site previously occupied by the flipped-out G27. **c.** Close-up view of the catalytic pocket in the SAH-bound G27-flipped-out state. The L1 loop is missing with the Y251 and F375 oriented away from the catalytic pocket, leaving the pocket accessible to the flipped-out G27. **d.** Comparison of low-pass filtered cryo-EM density maps of tRNA-Tyr in the SAH-bound G27-flipped-out (blue) and G27-unflipped (pink) states. The anticodon loop is well resolved in the G27-flipped-out state but becomes poorly resolved in the G27-unflipped state, suggesting increased conformational flexibility. **e.** Overall views of the TRMT1L-tRNA-Tyr complexes in the two SAH-bound states, showing differences in tRNA-Tyr conformation and engagement with the TRMT1L homodimer. tRNA-Tyr is shown as low-pass filtered density in blue (G27-flipped-out state) or pink (G27-unflipped state), whereas TRMT1L protomers A and B are shown as dark- and light-gray cartoons, respectively. **f-g.** Mutations of key residues (Y251 and F375) involved in gating the catalytic pocket impair TRMT1L-mediated m²,²G27 formation. In vitro transcribed tRNA-Tyr was incubated with purified Flag-TRMT1L WT or indicated mutant proteins and analyzed by immuno-Northern blotting with m²,²G antibody and densitometry quantification of the m²,²G signals. Data are presented as mean ± SD. N=3. **h-i.** The intact L1 loop is required for TRMT1L-mediated m²,²G27 formation. In vitro transcribed tRNA-Tyr was incubated with purified Flag-TRMT1L WT or the L1-loop mutant proteins and analyzed by immuno-Northern blotting with m²,²G antibody and densitometry quantification of the m²,²G signals. Data are presented as mean ± SD. N=3.

In the G27-flipped-out state, SAH occupies the open catalytic pocket, where it is stabilized by an extensive network of interactions involving the conserved residues D281, F283, M289, D307, L308, D357, D373 and F375 (Fig. 4b). To test whether these interactions are essential, we assessed the activity of mutants targeting some of these residues. Disruption of F283, D307, D373 and F375 substantially impaired methyltransferase activity, highlighting their importance in cofactor recognition and catalysis (Fig. 4c, d).

Adjacent to the SAH-binding pocket, the flipped-out G27 base sits in a shallow recess formed by G376, T400, and H460 (Fig. 4e). This arrangement places G27 close to the bound SAH and D373, which has been proposed to facilitate deprotonation of the exocyclic amine during methyl transfer ^23,27^. Both D373V and the isosteric D373N abolish activity (Fig. 4c, d); since asparagine preserves the size and hydrogen-bonding geometry that support the cofactor contact while removing the negative charge, the requirement at this position is for the carboxylate rather than for side-chain bulk or hydrogen bonding alone. To test the G27-binding recess directly, we mutated residues involved in G27 recognition. Mutating T400 to alanine impaired m²,²G27 formation (Fig. 4f, g). In contrast, H460A was as active as wild type. Extensive contact with the extruded base is therefore not in itself required for catalysis, and T400 may contribute to positioning G27 for methyl transfer rather than to retaining it. G376 lies next to the flipped-out base, and the G376I substitution, which introduces a bulky side chain that may sterically obstruct G27 positioning, abolished detectable activity (Fig. 4e-g). These results support a role for the G27-binding recess in positioning G27 for methylation.

Compared with the apo complex, the anticodon region and the G27-containing region adopt distinct conformations in the G27-flipped-out state and form additional contacts with both protomers (Fig. 4h). A36 flips out and contacts the adenine moiety of SAH and the side chain of L308 in protomer A, while G34 shifts and is sandwiched between R390 and R470 of protomer B. These contacts accompany a shift of the anticodon region toward protomer A, repositioning of the G27-containing region and a change in the conformation of the D-stem region that contains U20 (Fig. 4i). U35 lies in the anticodon loop engaged by protomer B, although density here is weaker than for the flanking nucleotides. Substitution of U35 with adenine or guanine reduced m²,²G27 formation to approximately one third of wild type, and the two substitutions were indistinguishable, indicating a requirement at this position that does not depend on which purine is present (Fig. 4j, k). Modeling a purine at position 35 suggests a potential steric clash with R387 that would arise only in the cofactor-bound state (Fig. 4l). These observations suggest that interactions between the anticodon region and the cofactor contribute to the tRNA rearrangements that favor G27 flipping toward the catalytic pocket.

### L1-loop stabilization and G27 repositioning

In the second conformation of the SAH-bound complex (Fig. 5a), the L1 loop adopts a stabilized conformation above the closed catalytic site, supported by an interaction network involving Y251, Q246, F375 and SAH (Fig. 5b). Y251 and F375 reorient toward SAH, contributing to closure of the pocket (Fig. 5b, c). G27 remains within the tRNA body, and the anticodon region has poorly resolved density and no longer contacts TRMT1L. Except for the T-stem and acceptor stem, which remain engaged, the rest of the tRNA is slightly displaced away from the protein, indicating a less engaged tRNA-binding conformation (Fig. 5d, e).

To relate L1-loop stabilization to tRNA positioning, we compared the two SAH-bound conformations at the catalytic site (Fig. 5b, c). The stabilized L1 loop occupies a position close to that of G27 when flipped out in the other conformation, suggesting that L1-loop stabilization may restrict G27 access to the catalytic pocket. In addition, L308, in a short turn after α3 helix, rearranges and no longer contacts the anticodon region, unlike in the G27-flipped-out state (Fig. 4h). Together, these observations suggest that L1-loop stabilization is associated with weakened anticodon-protein interactions and a less engaged tRNA-binding conformation in which G27 remains within the tRNA body.

The functional importance of the L1 loop is supported by mutagenesis. Disrupting the Y251-F375 interaction through Y251A or F375A abolished detectable methyltransferase activity (Fig. 5f, g). In contrast, Y251F, which removes only the hydroxyl group, and the Y251F/F375Y double substitution, which swaps the hydroxyl between the two positions, both retained activity (Fig. 5f, g). The requirement is therefore for the aromatic side chains rather than for the hydroxyl group. Replacing the TRMT1L L1 loop with the corresponding loop from TRMT1, which is five amino acids shorter, also abolished detectable activity (Fig. 5h, i). Together, these results indicate that both the interaction network and the length of the L1 loop are important for TRMT1L function.

## DISCUSSION

TRMT1L methylates a single nucleotide, G27, in a single tRNA species, yet our structures show it engaging much of that tRNA and, in the presence of SAH, extensively reshaping it: between these states the enzyme itself barely changes, while the substrate is rearranged. Earlier cellular studies ^16,19^ raised two questions that the structures reported here now make experimentally tractable: how TRMT1L restricts m²,²G27 formation to tyrosine tRNA, and how it supports modifications it does not itself catalyze.

The two protomers of the TRMT1L dimer have distinct roles. Protomer A carries the catalytic pocket, clamps the acceptor and T arms through its C-terminal region, and contacts the G27-containing region, whereas protomer B alone contacts the anticodon loop. Substituting the protomer B contacts abolishes m²,²G27 formation without preventing dimer assembly (Fig. 3h, i), and interface substitutions that weaken self-association also impair activity (Fig. 2g, h). Together with the contribution of the C-terminal zinc sites to dimer stability (Fig. 2j, k), these results support the dimer as a functional unit rather than an incidental feature of the purified protein.

Reorganization of the anticodon loop is a mechanistic feature of some tRNA-modifying enzymes, but in the cases we are aware of, it serves to expose a base within the loop for modification. G34 for tRNA-guanine transglycosylase ^28^, A34 for TadA ^29^, and G37 for TrmD and Trm5 ^30,31^. TRMT1L differs from these enzymes. In the cofactor-free complex, G27 remains base-paired and stacked at the first position of the anticodon stem even though the protomer B contacts are already formed, so initial anticodon engagement is not sufficient to drive flipping. In the SAH-bound state, G27 is flipped out toward the catalytic pocket, and the anticodon loop makes additional contacts with protomer A, including a contact between A36 and the cofactor (Fig. 4b, h). These contacts are functionally required: R390E/R470E abolishes m²,²G27 formation, and substituting U35 with adenine or guanine reduces it to about one third of wild type, consistent with a steric defect rather than loss of base-specific readout. The predicted clash between a purine at position 35 and R387 is a model-based prediction and not an observed contact (Fig. 4l). We therefore propose that cofactor-dependent reorganization of the anticodon region, accompanied by a change in the conformation of the D arm region that contains U20, delivers G27 to the catalytic pocket and couples exposure of the target base to the conformation of the tRNA as a whole. This proposal rests on a comparison of distinct structural states and does not establish the order or causality of these changes.

While this manuscript was in preparation, Gu et al. (2026) ^32^ reported, in a preprint, cryo-EM structures of dimeric TRMT1 and TRMT1L bound to tRNA. In TRMT1, G26 is already flipped in the apo complex, and cofactor then closes the L1 loop over the pocket. In their SAH-bound TRMT1L complex, G27 remains outside the catalytic pocket, which they interpret as a pre-catalytic state, and they concluded that the paralogs differ in their ability to remodel the target region. Our two SAH-bound states refine this view: in one, G27 remains stacked in the tRNA body, which may correspond to their structure, and in the other, G27 is extruded into the pocket, so TRMT1L can present its target base in association with cofactor-linked remodeling of the anticodon region. The paralogs may thus differ in how base extrusion is gated. Gu et al. ^32^ also report that TRMT1L-modified tRNA-Tyr accumulates m²G transiently before m²,²G, which is consistent with release between the two methylations, as our G27-unflipped state suggests.

More broadly, these structures suggest that TRMT1L achieves its narrow specificity not by recognizing G27 alone, but by docking tRNA-Tyr across a dimer and coupling cofactor binding to remodeling of the anticodon region and the adjacent D arm, so that presentation of the target base depends on the conformation of the tRNA as a whole. The same remodeling may also bear on the non-catalytic role of TRMT1L. TRMT1L supports acp³U20 formation by DTWD1 independently of its own catalytic activity ^16,19^, and in the SAH-bound state the D arm region that contains U20 adopts a distinct conformation (Fig. 4i). Together with the strand-annealing and dissociation activities described for fission yeast Trm1 ^21^, this raises the possibility that TRMT1L acts as a tRNA chaperone that prepares U20 for modification by DTWD1. Future studies will examine whether the remodeled tRNA is a preferred DTWD1 substrate.

## Data and code availability

Cryo-EM maps of the human TRMT1L-tRNA-Tyr complex in apo, SAH-bound G27-flipped-out and SAH-bound G27-unflipped states were deposited to the EMDataBank, with accessionnumbers EMDataBank: EMD-XXXXX, EMD-XXXXX, and EMD-XXXXX, respectively. The corresponding atomic models were deposited to the Protein Data Bank with accession numbers XXXX, XXXX, and XXXX.

## Acknowledgments

We thank the Structural Biology Imaging Center of the UTHealth McGovern Medical School for help with cryo-EM data collection. This work was supported by US National Institutes of Health grants R01 GM143587 (K.L.T.), and R01 GM158926 (C.D.).

## Author Contributions

T.L. and A.V. performed most of the experiments with the help of K.B., S-P.H., H-C.T and T-C.C. C.D., and K.LT. guided the research. T.L., A.V., C.D. and K.L.T. designed the experiments and wrote the paper with the input from all authors.

## Methods

### Cell culture and transfection

HEK 293T and parental HCT 116 cells were obtained from the American Type Culture Collection (ATCC, Manassas, VA, USA; CRL-3216 and CCL-247, respectively). Wild-type (WT) and TRMT1L-knockout (KO) HCT 116 cells were generated as previously described ^19^. All cells were cultured at 37°C in a 5% CO_2_ atmosphere in high-glucose Dulbecco’s modified Eagle’s medium (DMEM; HyClone SH30022.01) supplemented with 5% fetal bovine serum (FBS; HyClone SV30014.03), 100 U/ml penicillin, and 100 µg/ml streptomycin (HyClone SH40003.01). Cells were passaged using trypsin/EDTA before reaching 80% confluence and maintained in culture for no longer than 2 months after thawing. Cells were transfected with JetPRIME DNA and siRNA transfection reagent (Polyplus) at a ratio of 1:2 (reagent:DNA).

### Expression and purification of recombinant TRMT1L

The pFastBac1 plasmid encoding human TRMT1L with N-terminal 3xFLAG tag was transformed into DH10Bac competent cells. Bacmid DNA was isolated from white colonies, verified by PCR, and used to transfect Sf9 insect cells. Following three rounds of viral amplification, high-titer recombinant baculovirus was to infect High Five cells. At 48 h post-infection, 1 L of High Five cell culture was harvested, and the cells were lysed in buffer A (20 mM HEPES, pH 7.4, 500 mM NaCl, 10% glycerol, 2 mM MgCl2, 0.2 mM EDTA, 0.02% NP-40, 1 mM DTT and protease inhibitors) using a Dounce homogenizer. Cell lysate was clarified by centrifugation at 30,000 rpm for 40 min. The clarified supernatant containing TRMT1L was incubated with anti-FLAG M2 affinity beads for 2 h at 4℃. The beads were washed with buffer A for three times, followed by three washes with buffer B (20 mM HEPES, pH 7.4, 300 mM NaCl, 10% glycerol, 2 mM MgCl2, 0.2 mM EDTA, 0.02% NP-40, 1 mM DTT). TRMT1L was eluted with buffer B supplemented with 300 ng/µL FLAG peptide, and the eluates were analyzed by SDS-PAGE. The TRMT1L-containing eluates were concentrated and further purified by size-exclusion chromatography using a Superdex 200 Increase 10/300 GL column equilibrated with buffer C (20 mM HEPES, pH 7.4, 100 mM NaCl, 2 mM MgCl2, 1 mM DTT). Peak fractions corresponding to the TRMT1L dimer were pooled and concentrated.

### Preparation of TRMT1L-tRNA-Tyr and TRMT1L-tRNA-Tyr-SAH complexes for Cryo-EM

Purified TRMT1L was mixed with in vitro-transcribed tRNA-Tyr at a molar ratio of approximately 1: 2 and incubated on ice for 20 min. Following incubation, the mixture was centrifugated at 15,000 rpm for 10 min and subjected to size-exclusion chromatography using a Superdex 200 Increase 10/300 GL column equilibrated with buffer C (20 mM HEPES, pH 7.4, 100 mM NaCl, 2 mM MgCl2, 1 mM DTT). Eluted fractions were analyzed by SDS-PAGE with Coomassie blue staining for protein and by denaturing urea-PAGE with SYBR gold staining for RNA. Peak fractions containing the TRMT1L-tRNA-Tyr complex were pooled and concentrated to 1 mg/mL for cryo-EM specimen preparation. For grid preparation, 3 μL of the purified TRMT1L-tRNA-Tyr complex was applied to freshly glow-discharged R 0.6/1 UltraAuFoil@ Holey Gold grids. The grids were vitrified in liquid ethane using an FEI Vitrobot (Mark IV). For preparation of the TRMT1L-tRNA-Tyr-SAH complex, SAH was added to the concentrated TRMT1L-tRNA-Tyr complex to a final concentration of 250 µM. The sample was incubation for 10 min before application to cryo-EM grids and vitrification as described above.

### Cryo-EM data collection and processing

Single-particle cryo-EM images were collected using FEI Titan Krios (300 kV) electron microscope with a Gatan GIF Continuum K3 direct electron detector. The images were automatically acquired using EPU (Thermo Fisher Scientific) at underfocus values ranging from 0.8 to 2.0 µm with a pixel size of 0.85 Å px-1. Each micrograph was exposed for 2.9 s with a total does of about 58 e-Å-2, which was fractionated into 48 frames.

For Apo state, 5813 movie frames were aligned by MotionCor2. Gctf was used to estimate the parameters of contrast transfer function (CTF) for each image. The aligned micrographs were used to pick particles using Cryolo. Multiple rounds of 2D classification in Cryosparc were performed to remove junk particles and particles lacking clear features, resulting in a stack of 276,110 particles. A subset of 50,000 particles was used for 3D initial model generation using Cryosparc. An initial model showing TRMT1L forms a homodimer and one tRNA molecule binds to it. Multiple rounds of 3D classification were performed in RELION, resulting in 223,225 particles. 3D refinement in RELION produced a 3.7 Å-resolution map of the TRMT1L-tRNA complex. Next, the particles were subjected to Bayesian polishing in RELION, followed by heterogeneous refinement in cryoSPARC, which yielded 60,294 particles. These particles were subsequently subjected to non-uniform refinement, resulting in a map with a resolution of 3.22 Å.

For SAH bound state, 5,269 frames were aligned by MotionCor2. Gctf was used to estimate the parameters of contrast transfer function (CTF) for each image. The aligned micrographs were used to pick particles using Cryolo. Multiple rounds of 2D classification in Cryosparc were performed to remove junk particles, free tRNA, and particles lacking clear features, resulting in a stack of 394,524 particles. 3D refinement from these particles in RELION produced a 4.2 Å-resolution map of the TRMT1L-tRNA-SAH complex. Next, the particles were subjected to two rounds of 3D classification, yielding a dataset of 250,445 particles. These particles were subsequently subjected to 3D refinement and Bayesian polishing in RELION, followed by heterogeneous refinement and 3D classification in cryoSPARC. This processing resulted in two distinct maps: an SAH-bound, G27-unflipped state and an SAH-bound, G27-flipped-out state. The resolutions of final 3D maps were estimated using gold standard Fourier Shell correlation (FSC) curves with 0.143 criteria, and local resolutions were calculated using Cryosparc.

### Model building and refinement

To build the atomic model of the TRMT1L-tRNA complex in apo state, the AlphaFold 3 predicted structures of TRMT1L homodimer and tRNA-Tyr were used as templates and fitted into the density map of TRMT1L-tRNA apo state complex by rigid-body fitting in Chimera. The model was then manually built and adjusted in Coot, followed by real-space refinement in Phenix. The disordered regions in both TRMT1L subunits were not built because of missing or poor densities. To build the atomic model of the TRMT1L-tRNA-SAH complex, the apo state TRMT1L-tRNA atomic model was fitted into the density map of the TRMT1L-tRNA-SAH complex. Owing to the conformational change of tRNA, the tRNA was individually fitted into the map. The model was then manually built and adjusted in Coot, followed by real-space refinement in Phenix. The disordered regions in both TRMT1L subunits were not built because of missing or poor densities.

### Generation and expression of wildtype and TRMT1L mutants

PCR mutagenesis was performed using the Q5 Site-Directed Mutagenesis Kit (NEB E0554S) with primers listed in Table S1. Sequences of TRMT1L WT and mutant constructs were confirmed by Nanopore whole plasmid sequencing (Quintara Biosciences).

### Immunoprecipitation and immunoblotting

For co-immunoprecipitation experiments to assess TRMT1L dimer formation, cells were transfected with FLAG-TRMT1L WT ^19^ or mutants along with Myc-TRMT1L WT plasmids. 24h after transfection, cell were lysed in either low salt ELB buffer (50 mM HEPES, pH 7.2, 100 mM NaCl, 2 mM EDTA, 0.5% NP-40) or high salt ELB buffer (50 mM HEPES, pH 7.2, 250 mM NaCl, 2 mM EDTA, 0.5% NP-40) supplemented with protease inhibitors (aprotinin, Fisher BP250310; leupeptin, Fisher PI78435; AEBSF, Fisher BP635-500) on ice for 15 minutes. Lysates were then cleared by centrifugation at 22,000 × g for 10 minutes at 4°C, and the supernatant was incubated with 1 μg of FLAG or Myc (9E10; Santa Cruz Biotechnology sc-40) antibody for 1 hour with rotation at 4°C. Protein G Dynabeads (Invitrogen 10003D) were added for an additional 1 hour with rotation at 4°C. For immunoblotting, cells were lysed using RIPA buffer (25mM Tris-HCl pH 7.6, 150 mM NaCl, 1% NP-40, 1% Triton X-100, 1% sodium deoxycholate, and 0.1% SDS) and protease inhibitors on ice for 15 min. Lysates were cleared by centrifugation at 15,000 rpm for 10 minutes at 4°C. Protein concentrations were quantified using a BCA kit (Pierce 23225). Equal amounts of proteins were separated by SDS-PAGE and transferred to a 0.45 µm nitrocellulose membrane (Bio-Rad 1620115). The antibodies used for detection are: FLAG M2 (Sigma), Myc 9E10 (Santa Cruz Biotechnology sc-40), TRMT1L (Fortis Life Sciences A305-120A).

### Purification of Flag-tagged TRMT1L

Recombinant TRMT1L used for the in vitro m²,²G assays was purified as previously described with the following modifications ^19^. 5 µg of p3xFLAG-CMV-TRMT1L WT or mutant plasmid was transfected to HEK293T cells in a 10 cm dish. Two days after transfection, the cells were harvested and resuspended in lysis 0.75 ml buffer (20 mM HEPES (pH 7.5), 500 mM NaCl, 2 mMgCl_2_, 0.2 mM EDTA, 0.1 % NP40, 10% glycerol, 1 × protease inhibitor cocktail and 1 mM DDT) and lysed by sonication at intensity 4 with three cycles of 5 sec on-1 min off on ice. The lysates were cleared by centrifugation at 14,000 × *g* for 15 min at 4°C. The cleared lysate was mixed with 50 µl slurry of anti-DYKDDDDK Magnetic Agarose (Pierce A36797), and incubated at 4°C with rotation for 2 h. The agarose beads were washed three times 500 mM NaCl, 2 mMgCl_2_, 0.2 mM EDTA, 0.1 % NP40, 10% glycerol, and 1 mM DDT) followed by washing thrice with buffer B (300 mM NaCl, 2 mMgCl_2_, 0.2 mM EDTA, 10% glycerol, 1 × protease inhibitor cocktail and 1 mM DDT). For elution of the purified protein, the beads were mixed with 25 µl of 1.5 µg/µl 3x DYKDDDDK Peptide (Pierce A36805) in buffer B and incubated for 10 min at room temperature with rotation at 1400 rpm. The elution was repeated once for maximum recovery. The purified proteins were dialyzed on Slide-A-Lyzer MINI Dialysis Devices, 20K MWCO for FLAG peptide removal, in 100 ml buffer B, aliquoted and stored at -20°C. The purified proteins were resolved on SDS-PAGE gel and stained by SYPRO Ruby Protein Gel Stain (Invitrogen S12000).

### *In vitro* methyltransferase assay

In vitro methyltransferase assays were conducted using established methods ^19,26^. In brief, to prepare the *in vitro* transcribed tRNA, templates were amplified from double-stranded DNA gBlocks (IDT) containing a T7 promoter sequence upstream of the tRNA sequence. This amplification was performed by PCR using the primers listed in Table S1. The tRNA was transcribed *in vitro* from 50 ng of the PCR-amplified template using the MAXIscript™ T7 Transcription Kit (Invitrogen AM1312). The transcription reaction mixture was incubated at 37°C for 1 hour, followed by TURBO DNase (Invitrogen AM2239) treatment at 37°C for 15 min, as the manufacturer’s instructions suggest. The transcribed RNA was purified using the NEB Monarch RNA Cleanup kit (T2030L) and resolved on a 12% denaturing (7 M urea) polyacrylamide gel, stained with SYBR Gold (Invitrogen S11494) for visualization for quality control assessment. Prior to the *in vitro* methyltransferase reaction, the transcribed RNA was refolded. For refolding, the RNA was first denatured in 5 mM Tris (pH 7.5) and 0.16 mM EDTA at 95°C for 2 min, then immediately cooled on ice for 2 min. The denatured RNA was refolded in a buffer containing 50 mM HEPES, 3 mM MgCl_2_, and 50 mM NaCl. For the methyltransferase reaction, 100 ng of refolded RNA was mixed with 25 nM of purified protein in a reaction buffer containing 50 mM Tris (pH 7.5), 0.1 mM EDTA, 1 mM of freshly prepared DTT, and 0.5 mM *S*-adenosyl-methionine (SAM; NEB B9003S), and incubated at 30°C for 2h. Post-incubation, the RNA was denatured by mixing with 2x RNA loading dye and resolved on a 12% denaturing (8 M urea) polyacrylamide gel, followed by SYBR Gold staining and visualization under 302 nm UV transillumination. The RNA was then transferred to an Amersham Hybond N+ nylon membrane (GE Healthcare RPN303B) and crosslinked by exposure to 120 mJ/cm² UV. The m²,²G content in the crosslinked RNA was detected by immunoblotting with an m²,²G antibody (Abcam ab211488) ^26^.

### Quantification and statistical analysis

Data are shown as mean ± standard deviation. All data were generated from at least three biological replicates unless indicated. Each dot on dot-bar plot represents an independent biological replicate. A two-tailed Student’s t-test determined the difference between the two groups. *p* value < 0.05 were considered to be statistically significant. All the Western, immuno-Northern and Northern blots were quantified using Image J.

---

**Figure S1.**
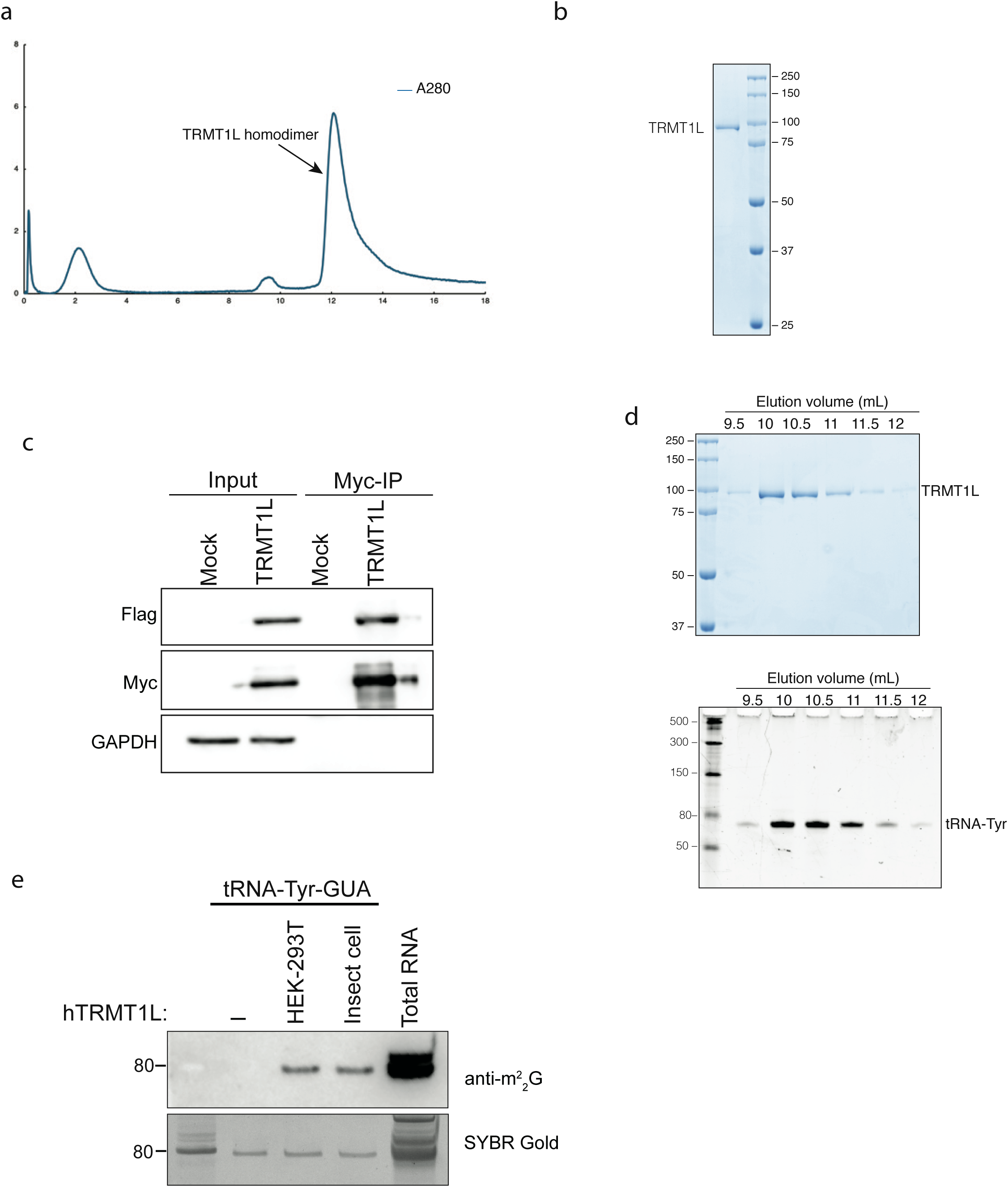
Biochemical and structural characterization of TRMT1L homodimerization. **a**. Size-exclusion chromatography profile of recombinant human TRMT1L showing a peak corresponding to the homodimeric assembly. **b.** Coomassie blue-stained SDS-PAGE gel of the TRMT1L peak fraction collected from the size-exclusion chromatography shown in a. **c.** Co-immunoprecipitation confirms TRMT1L homodimerization in cells. **d.** Coomassie blue-stained SDS-PAGE (top) and urea-TBE gel (bottom) analyses of peak fractions collected from the size-exclusion chromatography of the TRMT1L-tRNA-Tyr complex shown in Fig. 1b, confirming the co-elution of TRMT1L and tRNA-Tyr. **e.** Recombinant TRMT1L homodimer exhibits tRNA-Tyr methyltransferase activity.

**Figure S2.**
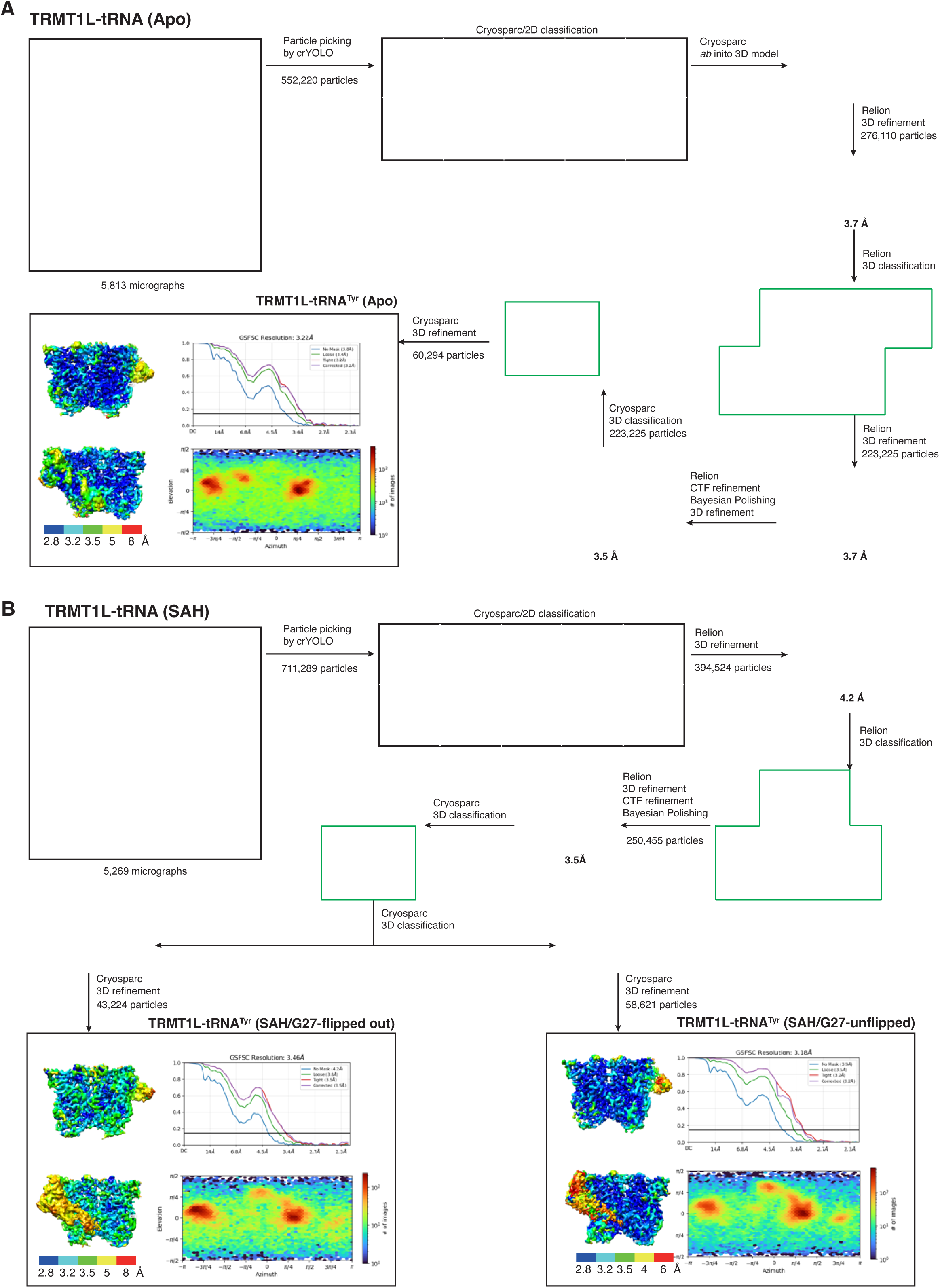
Cryo-EM data processing. **a.** Cryo-EM data processing workflow for the apo TRMT1L-tRNA-Tyr complex. **b.** Cryo-EM data processing workflow for the SAH-bound TRMT1L-tRNA-Tyr complexes.

**Figure S3.**
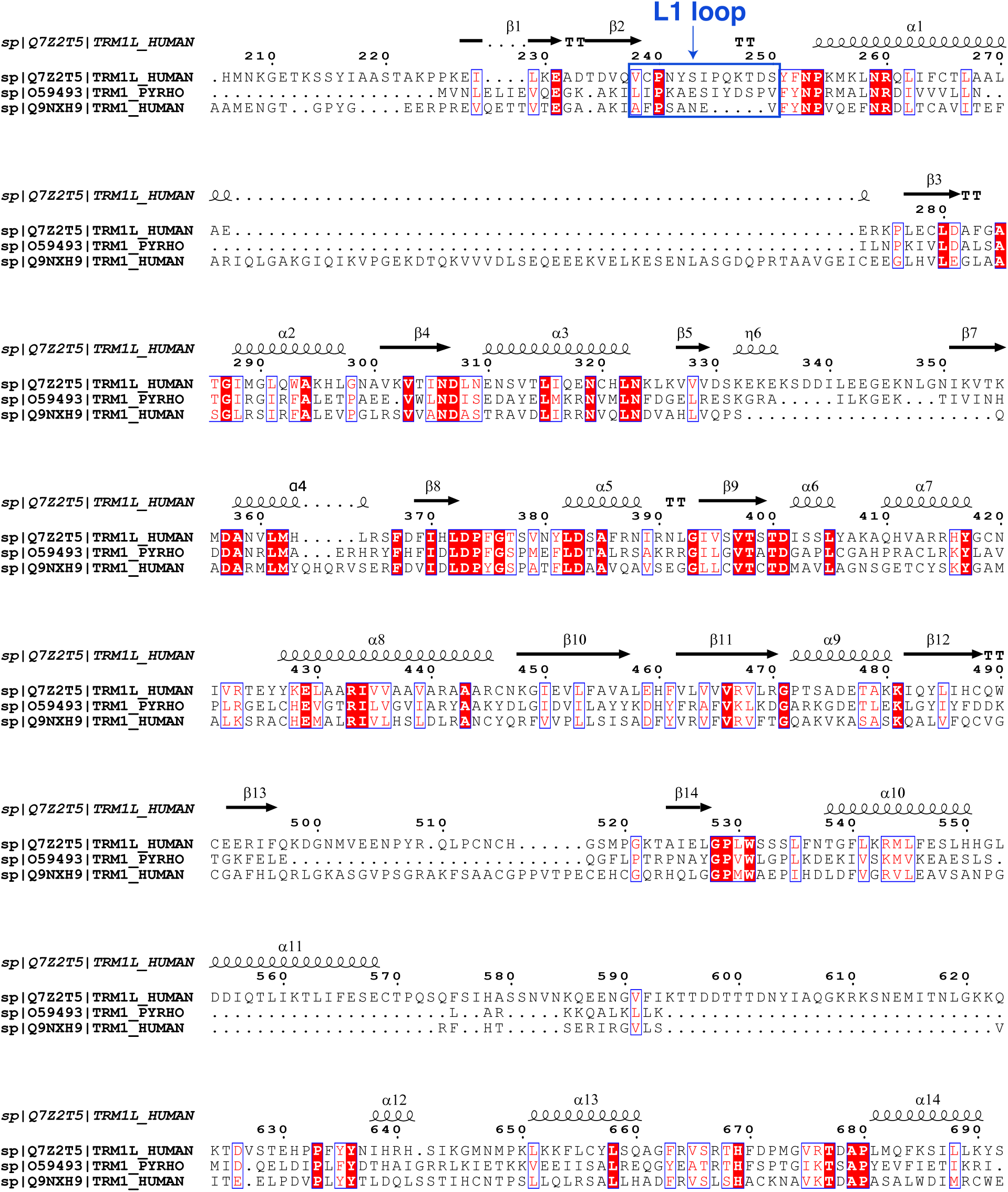
Multiple sequence alignment of the methyltransferase domains of human TRMT1L, human TRMT1 and Pyrococcus horikoshii Trm1. Multiple sequence alignment of human TRMT1L, P. horikoshii Trm1 and human TRMT1. Secondary-structure elements of human TRMT1L are shown above the alignment. Identical residues are highlighted in red, and similar residues are shown in red letters. The L1 loop region of TRMT1L and the corresponding regions in P. horikoshii Trm1 and TRMT1 are indicated by the blue box.

**Figure S4.**
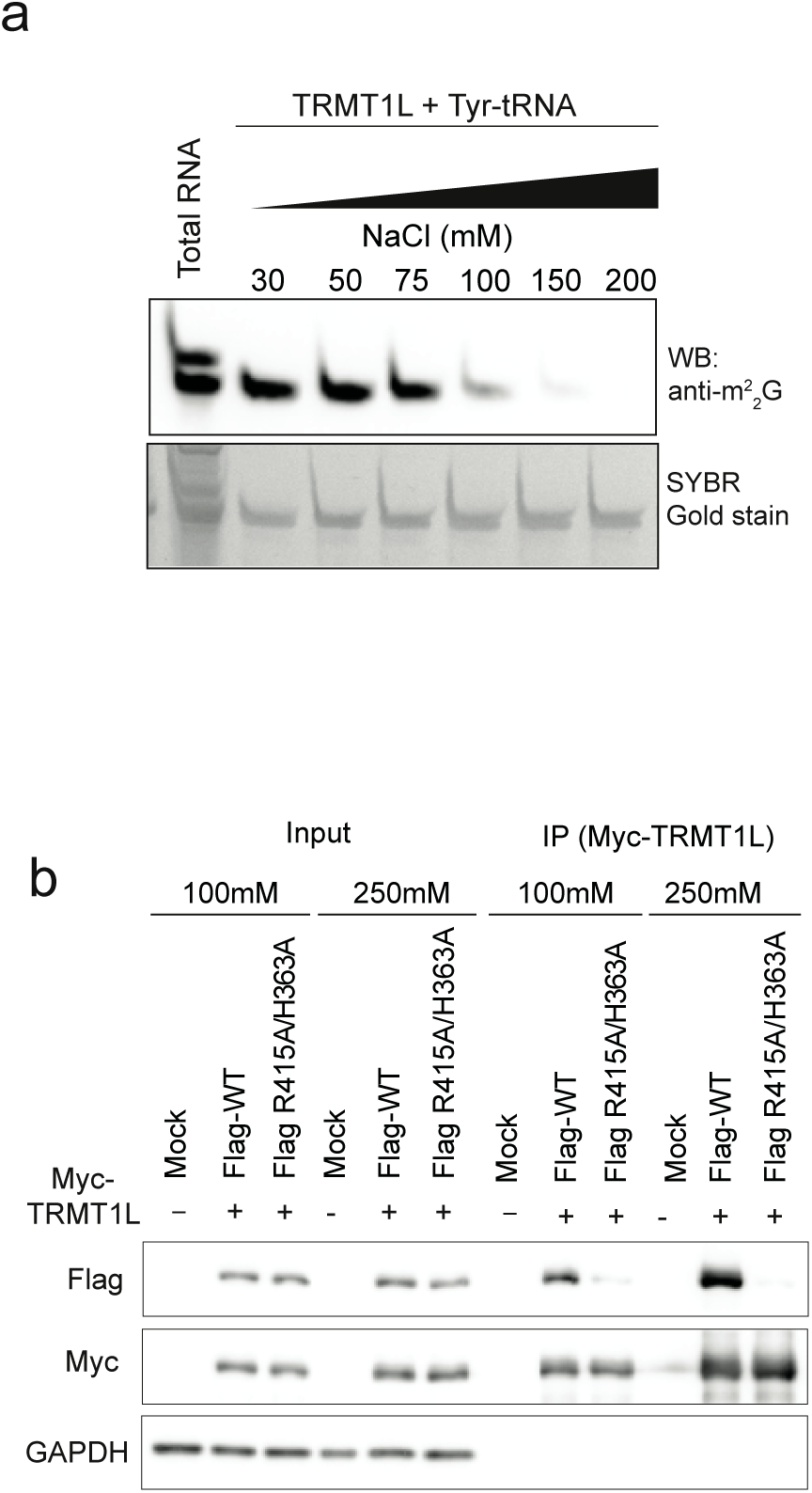
Effects of salt concentration and dimer-interface mutations on TRMT1L activity and dimerization **a.** Salt dependence of TRMT1L m²,²G27 methyltransferase activity on in vitro-transcribed tRNA-Tyr. **b.** Co-immunoprecipitation showing that the R415A/H363A double mutant impairs dimer formation at low salt.

**Figure S5.**
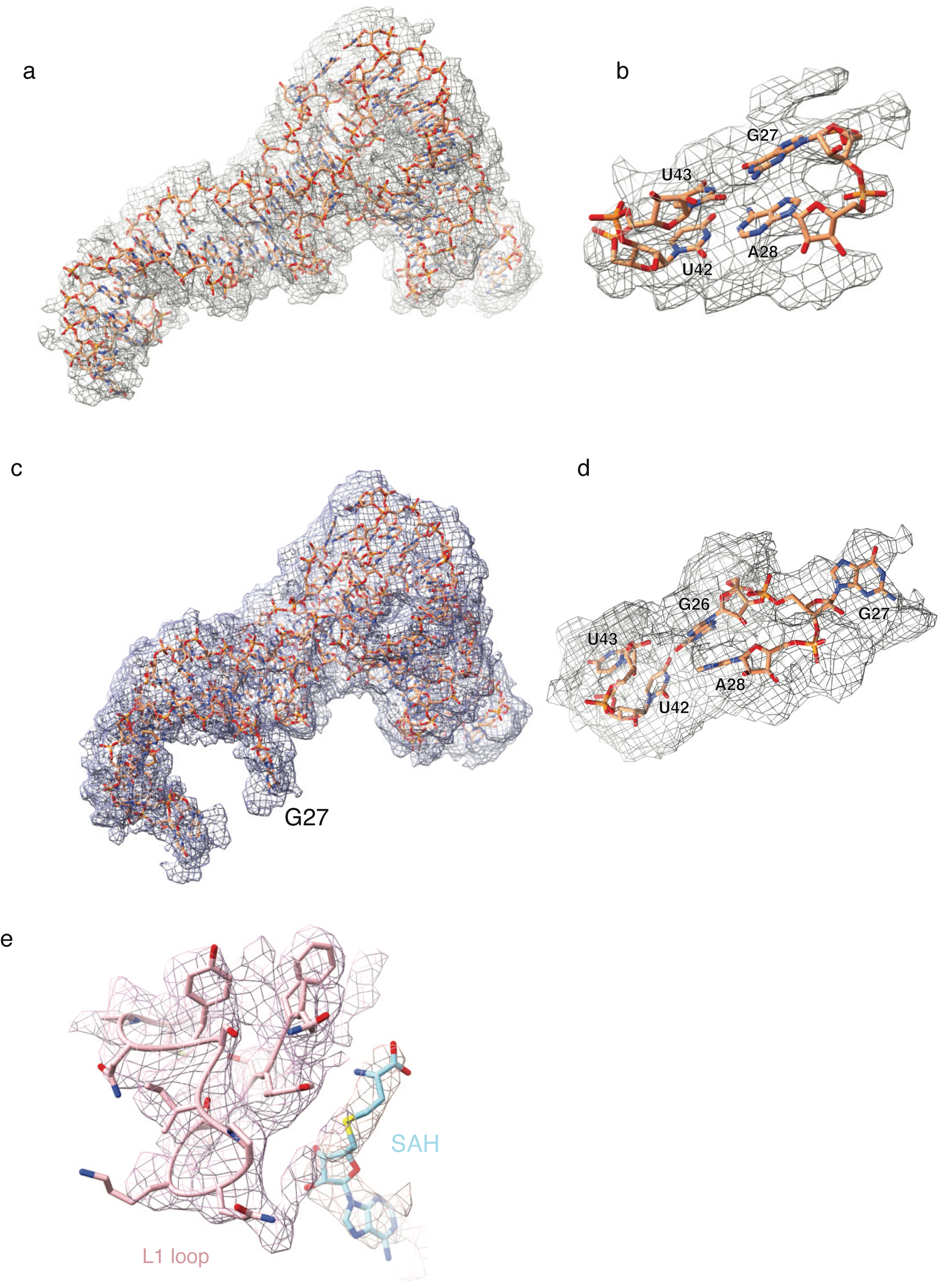
Cryo-EM density of tRNA-Tyr in the apo and SAH-bound states. **a.** Cryo-EM density of tRNA-Tyr in the apo TRMT1L-tRNA-Tyr complex. **b.** Close-up view of the G27 with cryo-EM density in the apo state. **c.** Cryo-EM density of tRNA-Tyr in the SAH-bound complex showing G27 base flipping. **d.** Close-up view of the G27 with cryo-EM density in the SAH-bound G27-flipped out state. **e.** Close-up view of the cryo-EM density corresponding to the stabilized L1 loop and bound SAH in the SAH-bound G27-unflipped state.

## References

1 Wei, J. & He, C. RNA modifications in gene regulation: Functions and pathways. Cell 189, 1591–1619 (2026). 10.1016/j.cell.2026.01.006

2 Suzuki, T. The expanding world of tRNA modifications and their disease relevance. Nat Rev Mol Cell Biol 22, 375–392 (2021). 10.1038/s41580-021-00342-0

3 Phizicky, E. M. & Hopper, A. K. The life and times of a tRNA. RNA 29, 898–957 (2023). 10.1261/rna.079620.123

4 Yared, M. J., Marcelot, A. & Barraud, P. Beyond the Anticodon: tRNA Core Modifications and Their Impact on Structure, Translation and Stress Adaptation. Genes (Basel) 15 (2024). 10.3390/genes15030374

5 Chan, C. T. et al. A quantitative systems approach reveals dynamic control of tRNA modifications during cellular stress. PLoS Genet 6, e1001247 (2010). 10.1371/journal.pgen.1001247

6 Dedon, P. C. & Begley, T. J. Dysfunctional tRNA reprogramming and codon-biased translation in cancer. Trends Mol Med 28, 964–978 (2022). 10.1016/j.molmed.2022.09.007

7 Zhang, W., Foo, M., Eren, A. M. & Pan, T. tRNA modification dynamics from individual organisms to metaepitranscriptomics of microbiomes. Mol Cell 82, 891–906 (2022). 10.1016/j.molcel.2021.12.007

8 Chujo, T. & Tomizawa, K. Human transfer RNA modopathies: diseases caused by aberrations in transfer RNA modifications. FEBS J 288, 7096–7122 (2021). 10.1111/febs.15736

9 Orellana, E. A., Siegal, E. & Gregory, R. I. tRNA dysregulation and disease. Nat Rev Genet 23, 651–664 (2022). 10.1038/s41576-022-00501-9

10 Burgess, R. W. & Storkebaum, E. tRNA Dysregulation in Neurodevelopmental and Neurodegenerative Diseases. Annu Rev Cell Dev Biol 39, 223–252 (2023). 10.1146/annurev-cellbio-021623-124009

11 Steinberg, S. & Cedergren, R. A correlation between N2-dimethylguanosine presence and alternate tRNA conformers. RNA 1, 886–891 (1995).

12 Urbonavicius, J., Armengaud, J. & Grosjean, H. Identity elements required for enzymatic formation of N2,N2-dimethylguanosine from N2-monomethylated derivative and its possible role in avoiding alternative conformations in archaeal tRNA. J Mol Biol 357, 387–399 (2006). 10.1016/j.jmb.2005.12.087

13 Pallan, P. S., Kreutz, C., Bosio, S., Micura, R. & Egli, M. Effects of N2,N2-dimethylguanosine on RNA structure and stability: crystal structure of an RNA duplex with tandem m2 2G:A pairs. RNA 14, 2125–2135 (2008). 10.1261/rna.1078508

14 Petrosyan, J. & Bohnsack, K. E. N2-methylguanosine and N2, N2-dimethylguanosine in cytosolic and mitochondrial tRNAs. Frontiers in RNA Research 2, 1460913 (2024). 10.3389/frnar.2024.1460913

15 Dewe, J. M., Fuller, B. L., Lentini, J. M., Kellner, S. M. & Fu, D. TRMT1-Catalyzed tRNA Modifications Are Required for Redox Homeostasis To Ensure Proper Cellular Proliferation and Oxidative Stress Survival. Mol Cell Biol 37 (2017). 10.1128/MCB.00214-17

16 Zhang, K. et al. Human TRMT1 and TRMT1L paralogs ensure the proper modification state, stability, and function of tRNAs. Cell Rep 44, 115092 (2025). 10.1016/j.celrep.2024.115092

17 Ellis, S. R., Morales, M. J., Li, J. M., Hopper, A. K. & Martin, N. C. Isolation and characterization of the TRM1 locus, a gene essential for the N2,N2-dimethylguanosine modification of both mitochondrial and cytoplasmic tRNA in Saccharomyces cerevisiae. J Biol Chem 261, 9703–9709 (1986).

18 Liu, J. & Straby, K. B. The human tRNA(m(2)(2)G(26))dimethyltransferase: functional expression and characterization of a cloned hTRM1 gene. Nucleic Acids Res 28, 3445–3451 (2000). 10.1093/nar/28.18.3445

19 Hwang, S. P. et al. TRMT1L-catalyzed m(2)(2)G27 on tyrosine tRNA is required for efficient mRNA translation and cell survival under oxidative stress. Cell Rep 44, 115167 (2025). 10.1016/j.celrep.2024.115167

20 Takakura, M., Ishiguro, K., Akichika, S., Miyauchi, K. & Suzuki, T. Biogenesis and functions of aminocarboxypropyluridine in tRNA. Nat Commun 10, 5542 (2019). 10.1038/s41467-019-13525-3

21 Porat, J. et al. Crosstalk between the tRNA methyltransferase Trm1 and RNA chaperone La influences eukaryotic tRNA maturation. J Biol Chem 299, 105326 (2023). 10.1016/j.jbc.2023.105326

22 Bujnicki, J. M., Leach, R. A., Debski, J. & Rychlewski, L. Bioinformatic analyses of the tRNA: (guanine 26, N2,N2)-dimethyltransferase (Trm1) family. J Mol Microbiol Biotechnol 4, 405–415 (2002).

23 Ihsanawati et al. Crystal structure of tRNA N2,N2-guanosine dimethyltransferase Trm1 from Pyrococcus horikoshii. J Mol Biol **383**, 871-884 (2008). 10.1016/j.jmb.2008.08.068

24 Awai, T. et al. Substrate tRNA recognition mechanism of a multisite-specific tRNA methyltransferase, Aquifex aeolicus Trm1, based on the X-ray crystal structure. J Biol Chem 286, 35236–35246 (2011). 10.1074/jbc.M111.253641

25 Constantinesco, F., Motorin, Y. & Grosjean, H. Characterisation and enzymatic properties of tRNA(guanine 26, N (2), N (2))-dimethyltransferase (Trm1p) from Pyrococcus furiosus. J Mol Biol 291, 375–392 (1999). 10.1006/jmbi.1999.2976

26 Hwang, S. P., Barondeau, K. & Denicourt, C. Protocol to detect m(2)(2)G modification on single tRNA iso-decoders by immuno-northern blotting. STAR Protoc 6, 103924 (2025). 10.1016/j.xpro.2025.103924

27 Awai, T. et al. Aquifex aeolicus tRNA (N2,N2-guanine)-dimethyltransferase (Trm1) catalyzes transfer of methyl groups not only to guanine 26 but also to guanine 27 in tRNA. J Biol Chem 284, 20467–20478 (2009). 10.1074/jbc.M109.020024

28 Xie, W., Liu, X. & Huang, R. H. Chemical trapping and crystal structure of a catalytic tRNA guanine transglycosylase covalent intermediate. Nat Struct Biol 10, 781–788 (2003). 10.1038/nsb976

29 Losey, H. C., Ruthenburg, A. J. & Verdine, G. L. Crystal structure of Staphylococcus aureus tRNA adenosine deaminase TadA in complex with RNA. Nat Struct Mol Biol 13, 153–159 (2006). 10.1038/nsmb1047

30 Ito, T. et al. Structural basis for methyl-donor-dependent and sequence-specific binding to tRNA substrates by knotted methyltransferase TrmD. Proc Natl Acad Sci U S A 112, E4197–4205 (2015). 10.1073/pnas.1422981112

31 Goto-Ito, S., Ito, T., Kuratani, M., Bessho, Y. & Yokoyama, S. Tertiary structure checkpoint at anticodon loop modification in tRNA functional maturation. Nat Struct Mol Biol 16, 1109–1115 (2009). 10.1038/nsmb.1653

32 Gu, X. et al. Molecular basis of site-selective N2-guanosine methylation in human tRNAs. Langtaosha (LTS) Preprints (2026). 10.65215/LTSpreprints.2026.10.09.000359

